## Supplementary material for "Water and chloride as allosteric inhibitors in WNK kinase osmosensing": Teixeira et al. Supplemental data

**This file includes:**

Supplemental Figure S1

Supplemental Tables S1-S9

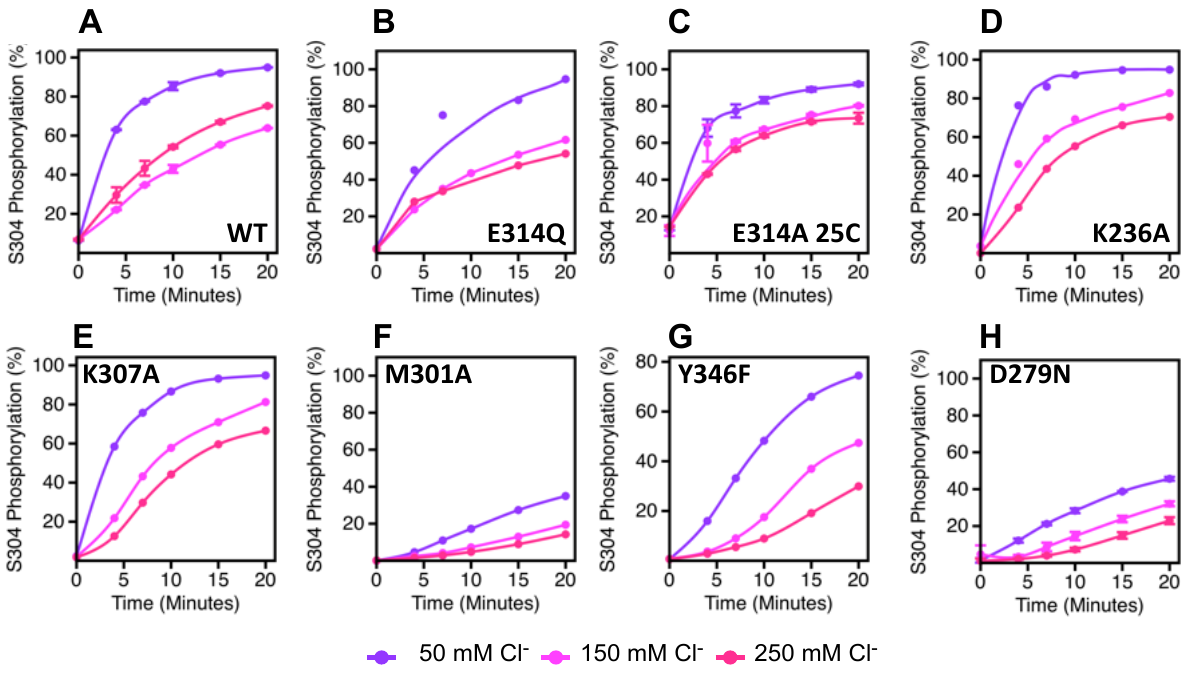

**Figure S1** Effects of chloride on uWNK3 AL-CL Cluster mutant autophosphorylation at S304. (A) Wild-type uWNK3 . (B) uWNK3/E314Q. (C) uWNK3/E214A at 25 ^o^C. (D) uWNK3/K236A (E) uWNK3/K307A (F) uWNK3/M301A. (G) uWNK3/Y346F. (H) uWNK3/D279N. Reactions run in 4 μM uWNK3, 30^o^C (unless otherwise indicated), at sodium chloride concentrations of 50 mM (purple), 150 mM (pink), and 250 mM (magenta). Bars indicate standard error from triplicate independent experiments.

**Table S1**. Crystallographic Data and Refinement of WNK1/S382A and WNK1/SA/PEG400

**WNK1/SA*	WNK1/SA/PEG400**

Space group	P1	P12_1_1

Unit cell dimensions *a, b, c* (Å)	38.25, 57.72, 65.60	38.32, 56.81, 65.28

Angles α, β, γ (°)	89.0, 89.6, 89.2	90, 95.4, 90

Wavelength (Å)	0.9795	0.9795

Resolution (Å)	50-2.0 (2.0-2.04)	30-2.0 (2.0-2.05)

Unique reflections (last shell)	38697	17297

Completeness (%) (last shell)	89 (83)	96 (93)

I/σ (last shell)	13.3(2.6)	29.8(8.8)

Rsym, Rpim (last shell)^a^	0.04, 0.045 (0.20, 0.24)	0.14, 0.06 (0.76, 0.29)

Redundancy (last shell)	1.7 (1.1)	7.6 (7.7)

CC1/2 (last shell)	0.99 (0.90)	0.56(0.55)

Wilson B factor	28.9	18.5

**Structure**

Rwork/Rfree^b^ (last shell)	0.16/0.22 (0.21/0.25)	0.199/0.233 (0.20/0.21)

Non-H protein atoms	4900	2281

Waters	453	250

Cryoprotectant (glycerol/PEG400) 9	0

RMSD in bond length (Å)^c^	0.006	0.016

RMSD in bond angles (°)^c^	1.38	1.65

Average B-values (Å^2^)	31.9	41.0

Ramachandran plot stats. (%)

Most favored region	94.6	90.8

Disallowed region	1.1	3.4

Molprobity Score	1.9	1.7

Residues missing from the model	None	379-385

^a^ R_sym_ = ∑ | I_avg_ - I_j_ | / ∑ I_j_.

^b^ R_factor_ = ∑ | F_o_ - F_c_ | / ∑ F_o_ , where F_o_ and F_c_ are observed and calculated structure factors,

respectively, R_free_ was calculated from a randomly chosen 5% of reflections excluded form

the refinement, and R_factor_ was calculated from the remaining 95% of reflections.

^c^ r.m.s.d is the root-mean-square deviation from ideal geometry.

* Newly crystallized form independent from 6CN9.

**Table S2.** Cell constant and Cα-Cα Superposition Comparisons

| A. Coordinate  sets | WNK1/SA  +PEG400 | WNK1/SA  +PEG400 | WNK1/SA  (6CN9) | pWNK1  (5W7T) |
| --- | --- | --- | --- | --- |
| Crystal system | Monoclinic  indexing | Triclinic  indexing | Triclinic | Monoclinic |
| Space Group | P2_1_ | P1 | P1 | P2_1_ |
| *a (*Å)  *b (*Å)  c *(*Å)  *α* (˚)  *β* (˚)  *γ* (˚) | 38.32  56.81  65.28  90.0  95.4  90.0 | 38.32  56.82  65.89  90.1  95.4  90.0 | 38.31  57.77  65.66  91.3  90.0  90.9 | 45.18  62.23  120.90  90.0  92.3  90.0 |
| Cα-Cα overlay | PEG-6CN9A | PEG-6CN9B | PEG-6CN9 | PEG-pWNK |
| *Å* | 0.79 | 1.3 | 3.2 | 0.82 |

**Table S3.** WNK3/1 expression level, and activation loop S308/S382 phosphorylation and activity

| **Mutants** | **Expression** | **Purification** | **Phos (%) from**  ***E. coli*** | **Auto-phos** | **Cl^-^ Sensitivity** |
| --- | --- | --- | --- | --- | --- |
| wt WNK3 | Very good | 26 mg/L | 98% | N/A | Very Sensitive |
| WNK3/E314A | Ok | 7 mg/L | 100% | Much faster | Insensitive |
| WNK3/E314Q | Ok | 6 mg/L | 99% | Faster | Insensitive |
| WNK3/K236A | Very good | 20 mg/L | 97% | Similar WT | Similar WT |
| WNK3/K307A | Good | 16 mg/L | 90% | Similar WT | Similar WT |
| WNK3/M301A | Very good | 23 mg/L | 96% | Slower | Very sensitive |
| WNK3/Y346F | Very good | 25 mg/L | 97% | Slower | Very sensitive |
| WNK3/D279N | Ok | 9 mg/L | 92% | Slower | Very sensitive |
| wt WNK1 | Good | 15 mg/L | 95% | Slower | Very Sensitive |
| WNK1/E388A | Ok | 10 mg/L | 94% | Faster WNK1 | < wt WNK1 |

**Table S4.** WNK1 and WNK3 peptides monitored by LC-MS

**Monitored peptide	(M+2H)^2+^	% acetonitrile**

Wild type WNK1	AKSVIGTPEFMAPEMY	885.5	22.5%

AKS*VIGTPEFMAPEMY	925.5	23.5%

WNK1/E388A	AKSVIGTPAFMAPEMY	885.5	18.0%

AKS*VIGTPAFMAPEMY	925.5	18.8%

Wild type WNK3	MRTSF	641.1	10.1%

MRTS*F	721.1	11.5%

AKSVIGTPEFMAPEMY	885.5	22.5%

AKS*VIGTPEFMAPEMY	925.5	23.5%

E314A	MRTSF	641.1	10.1%

MRTS*F	721.1	11.5%

AKSVIGTPAFM	561.5	18.0%

AKS*VIGTPAFM	601.5	18.8%

K236A	MRTSF	641.1	10.1%

MRTS*F	721.1	11.5%

AKSVIGTPEFMAPEMY	885.5	22.5%

AKS*VIGTPEFMAPEMY	925.5	23.5%

K307A	MRTSF	641.1	10.1%

MRTS*F	721.1	11.5%

AASVIGTPEFMAPEMY	856.9	30.3%

AAS*VIGTPEFMAPEMY	897.0	33.2%

M301A	ARTSF	581.1	7.3%

ARTS*F	661.1	8.6%

AKSVIGTPEFMAPEMY	885.5	22.5%

AKS*VIGTPEFMAPEMY	925.5	23.5%

Y346F	MRTSF	641.1	10.1%

MRTS*F	721.1	11.5%

AKSVIGTPEFMAPEMY	885.5	22.5%

AKS*VIGTPEFMAPEMY	925.5	23.5%

D297N	MRTSF	641.1	10.1%

MRTS*F	721.1	11.5%

AKSVIGTPEFMAPEMY	885.5	22.5%

AKS*VIGTPEFMAPEMY	925.5	23.5%

______________________________________________________________________________________

Wild type and mutant WNK1- and WNK3-specific chymotrypsin-derived activation loop peptides monitored

to generate autophosphorlation progress curves. M*/z* for the (M+2H)^2+^ of peptide and percent

acetonitrile where peptide elutes from a C18 HPLC column (0.1% formic acid in acetonitrile/water

HPLC mobile phase).

**Table S5.** uWNK1 and uWNK3 autophosphorylation assay conditions

| Component |  |  |  |
| --- | --- | --- | --- |
| HEPES pH7.4 | 20 mM | 20 mM | 20 mM |
| Mg Gluconate | 20 mM | 20 mM | 20 mM |
| Final [Cl^-^] | 50 mM | 150 mM | 250 mM |
| ATP | 5 mM | 5 mM | 5 mM |
| uWNK1/uWNK3 | 4 μM | 4 μM | 4 μM |
| Final Volume | 50 μL | 50 μL | 50 μL |
| Guanidine-HCl to final conc. of 1M to stop the reaction.. | | |  |

**Table S6** Wildtype and E388A uWNK1 autophosphorylation were fit to a basic autocatalytic mechanism using DynaFit software. Modeled progress curves superimposed on mass spectrometry data are shown in Figure 5E and 5F.

**Mechanism**

uWNK + pWNK →	2 pWNK	:k_P_

uWNK + ion	⇄	uWNK_ion	:ion_on	ion_off

**ODE System**

d[uWNK]/dt	=	- kP[uWNK][pWNK] - ion_on[uWNK][ion] + ion_off[uWNK_ion]

d[pWNK]/dt	=	+ kP[uWNK][pWNK]

d[ion]/dt	=	- ion_on[uWNK][ion] + ion_off[uWNK_ion]

d[uWNK_ion]/dt	=	+ ion_on[uWNK][ion] - ion_off[uWNK_ion]

**Optimized Parameters**

**uWNK1 wt	uWNK1 E388A**

| Parameter | Initial Value | Final Value | Std. Error | CV(%) |
| --- | --- | --- | --- | --- |
| KP | 0.18 | 0.183 | 0.008 | 4.4 |
| Ion_on | 2e6 | 3.54e^-6^ | 5.4e^-7^ | 15.2 |
| Ion_off | 0.2 | 0.119 | 0.013 | 11.1 |
| Kionbound |  | 2.98e^-5^ |  |  |

| Parameter | Initial Value | Final Value | Std. Error | CV(%) |
| --- | --- | --- | --- | --- |
| KP | 0.18 | 0.150 | 0.007 | 4.7 |
| Ion_on | 2e6 | 4.15e^-6^ | 6e^-7^ | 14.4 |
| Ion_off | 0.2 | 0.103 | 0.011 | 10.5 |
| Kionbound |  | 4.03e^-5^ |  |  |

**Table S7.** Molecular weight vs [WNK] by static light scattering

| **WT/ Mutants** | **MW at 0.8 mg/mL (KDa)** | **St. Dev.** | **MW at 1.8 mg/mL** | **St. Dev.** |
| --- | --- | --- | --- | --- |
| wt WNK3 | 65 | 0.6 | 78 | 0.5 |
| WNK3/E314Q | 51 | 0.2 | 52 | 0.1 |
| WNK3/E314A | 56 | 0.1 | 77 | 1.1 |
| WNK3/K236A | 61 | 0.3 | 85 | 0.2 |
| WNK3/K307A | 59 | 0.3 | 65 | 0.3 |
| WNK3/M301A | 64 | 0.4 | 69 | 0.2 |
| WNK3/Y346F | 68 | 0.8 | 76 | 0.1 |
| WNK3/D279N | 39 | 0.1 | 41 | 0.1 |
| wt WNK1 | 38 | 0.5 | 35 | 0.1 |
| WNK1/E388A | 39 | 0.1 | 35 | 0.6 |

**Table S8**. Crystallographic Data and Refinement of WNK3/SA/E314A

**WNK3/SA/E314A**

Space group	P12_1_1

Unit cell dimensions *a, b, c* (Å)	50.17, 113.60, 67.52

Angles α, β, γ (°)	90, 101.4, 90

Wavelength (Å)	0.9795

Resolution (Å)	43-3.3 (3.36-3.3)

Unique reflections (last shell)	7500

Completeness (%) (last shell)	70(55)

I/σ (last shell)	9.8(1.0)

Rsym, Rpim (last shell)^a^	0.17, 0.08 (0.76, 0.54)

Redundancy (last shell)	5.2 (3.7)

CC1/2 (last shell)	0.96(0.60)

Wilson B factor	81.9

**Structure**

Rwork/Rfree^b^ (last shell)	0.188/.270 (0.30/0.42)

Non-H protein atoms	4255

Waters	117

RMSD in bond length (Å)^c^	0.004

RMSD in bond angles (°)^c^	1.62

Average B-values (Å^2^)	87.1

Ramachandran plot stats. (%)

Most favored region	88.4

Disallowed region	1.6

Molprobity Score	1.9

Residues missing from the model	A:/303-314;B:/303-315

^a^ R_sym_ = ∑ | I_avg_ - I_j_ | / ∑ I_j_.

^b^ R_factor_ = ∑ | F_o_ - F_c_ | / ∑ F_o_ , where F_o_ and F_c_ are observed and calculated structure factors,

respectively, R_free_ was calculated from a randomly chosen 5% of reflections excluded form

the refinement, and R_factor_ was calculated from the remaining 95% of reflections.

^c^ r.m.s.d is the root-mean-square deviation from ideal geometry.

**Table S9** Key resources.

| Reagent type (species)  or resource | Designation | Source or reference | Identifiers | Additional information |
| --- | --- | --- | --- | --- |
| Recombinant DNA reagent | WNK1(194-483) (plasmid) | This paper |  | GenScript |
| Recombinant DNA reagent | WNK1(194-483)_E388A (plasmid) | This paper |  | GenScript |
| Recombinant DNA reagent | WNK3(118-409) (plasmid) | (Akella *et al*, 2021) |  | GenScript |
| Recombinant DNA reagent | WNK3(118-409)_K236A (plasmid) | This paper |  | GenScript |
| Recombinant DNA reagent | WNK3(118-409)_D279N (plasmid) | This paper |  | GenScript |
| Recombinant DNA reagent | WNK3(118-409)_M301A (plasmid) | This paper |  | GenScript |
| Recombinant DNA reagent | WNK3(118-409)_K307A (plasmid) | This paper |  | GenScript |
| Recombinant DNA reagent | WNK3(118-409)_E314Q (plasmid) | This paper |  | GenScript |
| Recombinant DNA reagent | WNK3(118-409)_E314A (plasmid) | This paper |  | GenScript |
| Recombinant DNA reagent | WNK3(118-409)_Y346F (plasmid) | This paper |  | GenScript |
| Recombinant DNA reagent | WNK3(118-409)_S308A_E314A (plasmid) | This paper |  | GenScript |
| Recombinant DNA reagent | PP1cϒ (plasmid) | (Barford & Keller, 1994) |  | From Depaoli-Roach |
| Peptide, recombinant protein | GST-OSR1(314-344) | (Taylor *et al*, 2018) |  | From Melanie Cobb |
| Peptide, recombinant protein | Lambda phosphatase | Santa Cruz Biotechnology | Cat# sc-200312 |  |
| Commercial assay or kit | ADP-Glo Max Assay | Promega | Cat# V7001 |  |

Akella R, Humphreys JM, Sekulski K, He H, Durbacz M, Chakravarthy S, Liwocha J, Mohammed ZJ, Brautigam CA, Goldsmith EJ (2021) Osmosensing by WNK Kinases. *Mol Biol Cell* 32: 1614-1623

Barford D, Keller JC (1994) Co-crystallization of the catalytic subunit of the serine/threonine specific protein phosphatase 1 from human in complex with microcystin LR. *J Mol Biol* 235: 763-766

Taylor CAt, An SW, Kankanamalage SG, Stippec S, Earnest S, Trivedi AT, Yang JZ, Mirzaei H, Huang CL, Cobb MH (2018) OSR1 regulates a subset of inward rectifier potassium channels via a binding motif variant. *Proc Natl Acad Sci U S A* 115: 3840-3845
